# The value of time in the invigoration of human movements when interacting with a robotic exoskeleton

**DOI:** 10.1101/2023.03.21.533648

**Authors:** Dorian Verdel, Olivier Bruneau, Guillaume Sahm, Nicolas Vignais, Bastien Berret

## Abstract

Time and effort are critical factors that are thought to be subjectively balanced during the planning of goal-directed actions, thereby setting the vigor of volitional movements. Theoretical models predicted that the value of time should then amount to relatively high levels of effort. However, the time-effort tradeoff has so far only been studied for a narrow range of efforts. Therefore, the extent to which humans can invest in a time-saving effort remains largely unknown. To address this issue, we used a robotic exoskeleton which significantly varied the energetic cost associated with a certain vigor during reaching movements. In this situation, minimizing the time-effort tradeoff would lead to high and low human efforts for upward and downward movements respectively. Consistent with this prediction, results showed that all participants expended substantial amounts of energy to pull on the exoskeleton during upward movements and remained essentially inactive by harnessing the work of gravity to push on the exoskeleton during downward movements, while saving time in both cases. These findings show that a common tradeoff between time and effort can determine the vigor of reaching movements for a wide range of efforts, with time cost playing a pivotal role.

## 1 Introduction

Most actions in daily life require us to select the speed or duration of goal-directed movements, that is, their *vigor* [1]. Thus, it is an ubiquitous feature of volitional actions, the setting of which is thought to be rooted in the basal ganglia [2, 3], in particular the striatum [4–10]. Current works suggest that vigor essentially reflects the internal value, or utility, of a given action [1,11, 13]. Numerous behavioral studies have shown that vigor is indeed modulated by the expected reward of the task at hand [14–19], with reward tending to be discounted over time [20–22]. However, if the modulation of vigor allows to modify the time needed to accomplish a task, it also affects the energy expenditure. Interestingly, reward has also been found to increase the propensity to put extra effort into a task [11, 23]. Therefore, movement vigor may generally result from the maximization of a capture rate, such as the sum of all rewards achieved minus all efforts expended, divided by the time. This global tendency has been observed in humans and many other species in foraging-like tasks [24–27]. An alternative formulation considers vigor as the outcome of the minimization of a subjective weighting between a cost of time (CoT) and a cost of movement, modulated by the expected reward [20, 21], which is convenient to model vigor in reaching tasks [28, 29]. When reward is not explicit (e.g., pointing to a light spot), movement vigor could then be determined by a common tradeoff between time and effort, which could represent a trait-like feature of individuality [30–34]. Empirical evidence of such a subjective CoT was recently reported in an isometric reaching task without explicit reward [35]. Based on this premise, several computational models were developed to account for the vigor of individuals during walking [34] and reaching [20,29,31,35,36], from a similar minimum time-effort (MTE) principle. Estimation of the underlying CoT in reaching was obtained from point-to-point movements of various amplitudes, using effort costs traditionally represented in motor control [29,31], even though other factors such as accuracy or comfort may also modulate vigor in general [37–41].

Interestingly, computational models revealed that the putative CoT should actually grow quickly to account for the vigor of self-paced pointing movements, such that time could amount to relatively high levels of effort. In other words, people could be prone to expend substantial energy to avoid excessively long movement times. Previous paradigms did not allow to test this prediction because the energetic cost of actions was too small or varied marginally through the different conditions of the task [11, 20, 27, 31, 35, 36]. Furthermore, while moving faster requires more energy expenditure, it does not necessarily have to come from human muscles, as demonstrated by using an electric bike or cycling downhill for instance. Therefore, do people rely on a common time-effort tradeoff to set movement vigor when the effort term is broadly varied experimentally?

Here, we designed an original experiment leveraging the versatility of a robotic exoskeleton to investigate this question. Two conditions requiring either a high or low energy expenditure to move with a similar vigor were implemented. The task consisted of performing vertical forearm movements to point-light targets while wearing the exoskeleton (Fig. 1A). During upward movements, the exoskeleton provided an assistance of duration *T_j_* along a predefined human-like trajectory so that the participant could comfortably and accurately complete the task without any effort. Crucially, this duration could be significantly longer than the participant’s preferred movement duration in the task, *T*_*h*,0_. In this case, the MTE theory predicts that all participants should be prone to energize the movement by pulling on the exoskeleton (Fig. 1B). To induce high levels of effort, and strongly penalize potential time savings, the robot applied a viscous-like resistance proportional to the participants’ maximum voluntary force as soon as they outpaced it. During downward movements, we took advantage of gravity to design a different assistance whereby saving a similar amount of time as for upward movements would instead require virtually no effort. In this case, the MTE theory predicts that all participants should remain practically inactive to behave optimally (Fig. 1C). This apparatus allowed for a significant departure from the MTE predictions depending on the participants’ choices. For instance, participants could choose to remain inactive in all conditions, thus failing to save time when relevant in the sense of the MTE theory. In contrast, participants could actively put energy into the task in all conditions, thus failing to save effort when relevant in the sense of the MTE theory. Thus, the results will determine whether vigor is the result of a common time-effort tradeoff during reaching movements whose energy cost for a certain duration varies greatly, or whether the MTE principle should be revised.

**Figure 1:**
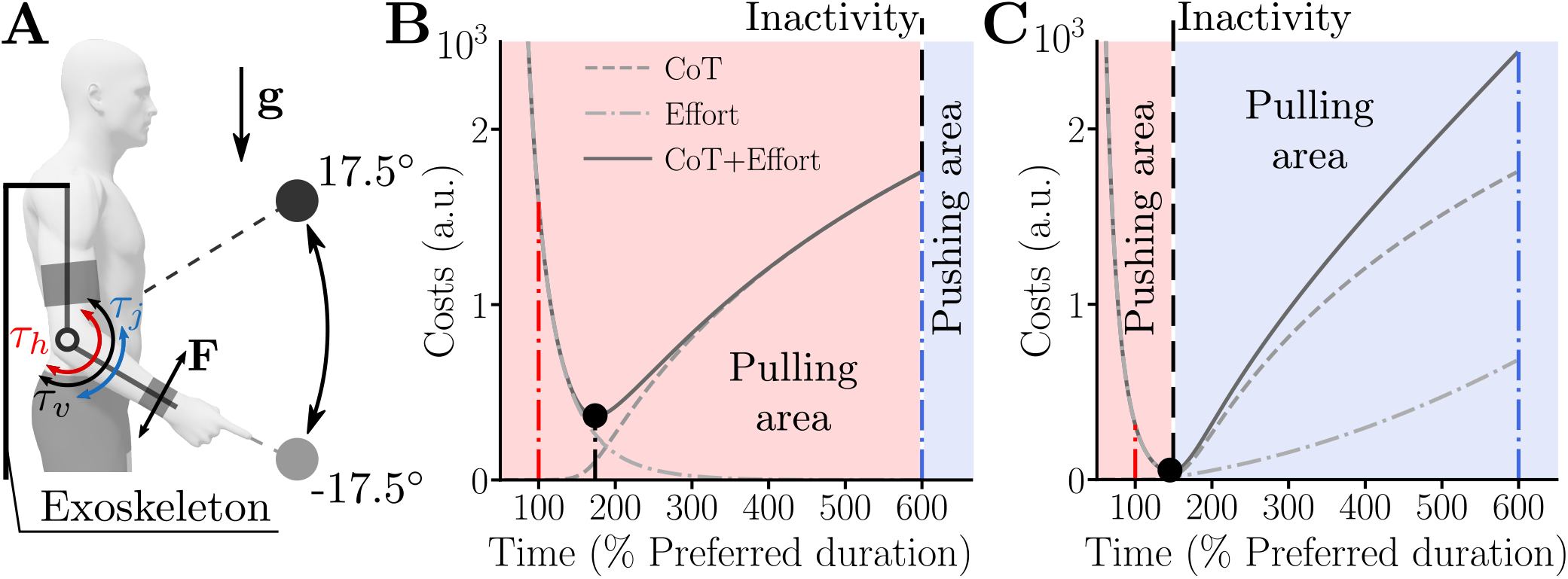
Illustration of theoretical predictions for an assistance 6x slower than the nominal vigor of the participant in the task (*i.e., T_j_* = 600%*T*_*h*,0_). **A.** Different input torques involved in the task. The term *τ_j_* is the torque provided by the robot as a biological movement assistance (here minimum jerk trajectory of duration *T_j_*) as long as the participant does not outpace the planned trajectory. The term *τ_h_* is the net torque produced by the human muscles (*τ_h_* = 0 when the participant is inactive). The term “pulling” (respectively “pushing”) refers to a participant generating an upward/positive (respectively downward/negative) torque *τ_h_*. The term *τ_v_* is a viscous-like torque applied by the robot, which is replacing *τ_j_* as soon as the participant choose to outpace the planned trajectory. F is the measured interaction force. **B**. Possible strategies during upward movements in terms of effort and time costs (see Equation 8 for details regarding the cost function). The red vertical line highlights the costs associated with the participant’s preferred duration *T*_*h*,0_. The blue vertical line highlights the costs associated with the exoskeleton’s planned duration. The black disk represents the optimal strategy in the sense of a MTE tradeoff. During upward movements, the participant could only save time by actively pulling on the exoskeleton (i.e., *τ_h_* > 0), which is represented by the shaded red area. The participant could also remain inactive and be moved by the robot, which is represented by the vertical dashed black line (inactivity). Otherwise, the participant could actively push against the exoskeleton (i.e., *τ_h_* < 0), although it would mean voluntarily wasting both time and effort. **C**. Possible strategies during downward movements in terms of effort and time costs. The pulling (shaded blue) and pushing (shaded red) areas are different from panel **B** because both pushing and pulling can allow to save time in this condition (although the latter strategy would be non-optimal from the MTE perspective). The critical difference for downward movements is that participants could save time by simply dropping their forearm, thereby passively pushing on the exoskeleton thanks to their own weight (dashed black line labelled inactivity). A strong deviation from this nearly optimal strategy could be observed if participants use a fixed effort-based heuristic to save time, by either actively pulling (to compensate for a part of the weight) or actively pushing on the exoskeleton.

## 2 Results

In this experiment, we asked *N* =12 participants to perform reaching movements to point-light targets at their preferred pace. The movements consisted of a discrete sequence of vertical elbow flexions and extensions. Both the target and a visual feedback of the participant’s current position were displayed on a large screen in front of the participant. Our experiment was divided in two sessions. In the first session *(baseline),* the exoskeleton was controlled in transparent mode, that is, no assistance was provided by the robot that compensated for its own dynamics and minimized interaction efforts [42, 43]. In this session, before being installed in the exoskeleton, the participants also performed a maximum isometric voluntary force (MVF) test, performed using an 1-axis force transducer (the reader is deferred to the Methods section for details regarding the procedure). The main objectives of the baseline session were to estimate the nominal vigor of the participants (*i.e.*, their preferred movement duration in the task) and their maximal force characteristics. This allowed to design a subject-specific assistance, which was normalized with respect to time and effort for the subsequent test session. Knowing the nominal vigor of the participants in the task further allowed us to infer the CoT for the optimal control simulations [29, 35]. In the second session *(test),* we asked the same participants to perform similar movements but with a personalized assistance provided by the robot. To this aim, we programmed the exoskeleton to follow minimum jerk trajectories of different durations, ranging from the participant’s preferred vigor (*T_j_* = 100%*T*_*h*,0_) to a 6x slower vigor (*T_j_* = 600%*T*_*h*,0_). Participants could decide to outpace the planned trajectory at any time during the movement. For upward movements, this required an active effort from the participant but, for downward movements, the planned trajectory could be outpaced by simply remaining inactive due to the effects of gravity. For both movement directions, when the planned trajectory was outpaced, the robot applied a viscous-like resistance proportional to the participant’s MVF (see Equation 3). The reader is deferred to the Materials and Methods section for more details about all the procedures.

### 2.1 Baseline session

In the baseline session, participants performed self-paced vertical pointing movements of four different amplitudes without active assistance/resistance from the robot. Qualitatively, the velocity profiles were overall bell-shaped as it is commonly observed for unrestrained point-to-point movements of this type (see Figures 2A,B,D,E). The only exception was for the largest movement amplitude which tended to exhibit a correction near the end of the movement (see Figures 2B,E). Importantly for our purpose, we observed the classical affine amplitude-duration relationship that characterizes the vigor of self-paced reaching movements [31, 33, 44, 45]. This relationship was observed at both individual and population levels, for upward and downward movements separately (see Figures 2C,F). These findings are consistent with results from previous studies with the same exoskeleton [42, 46].

**Figure 2:**
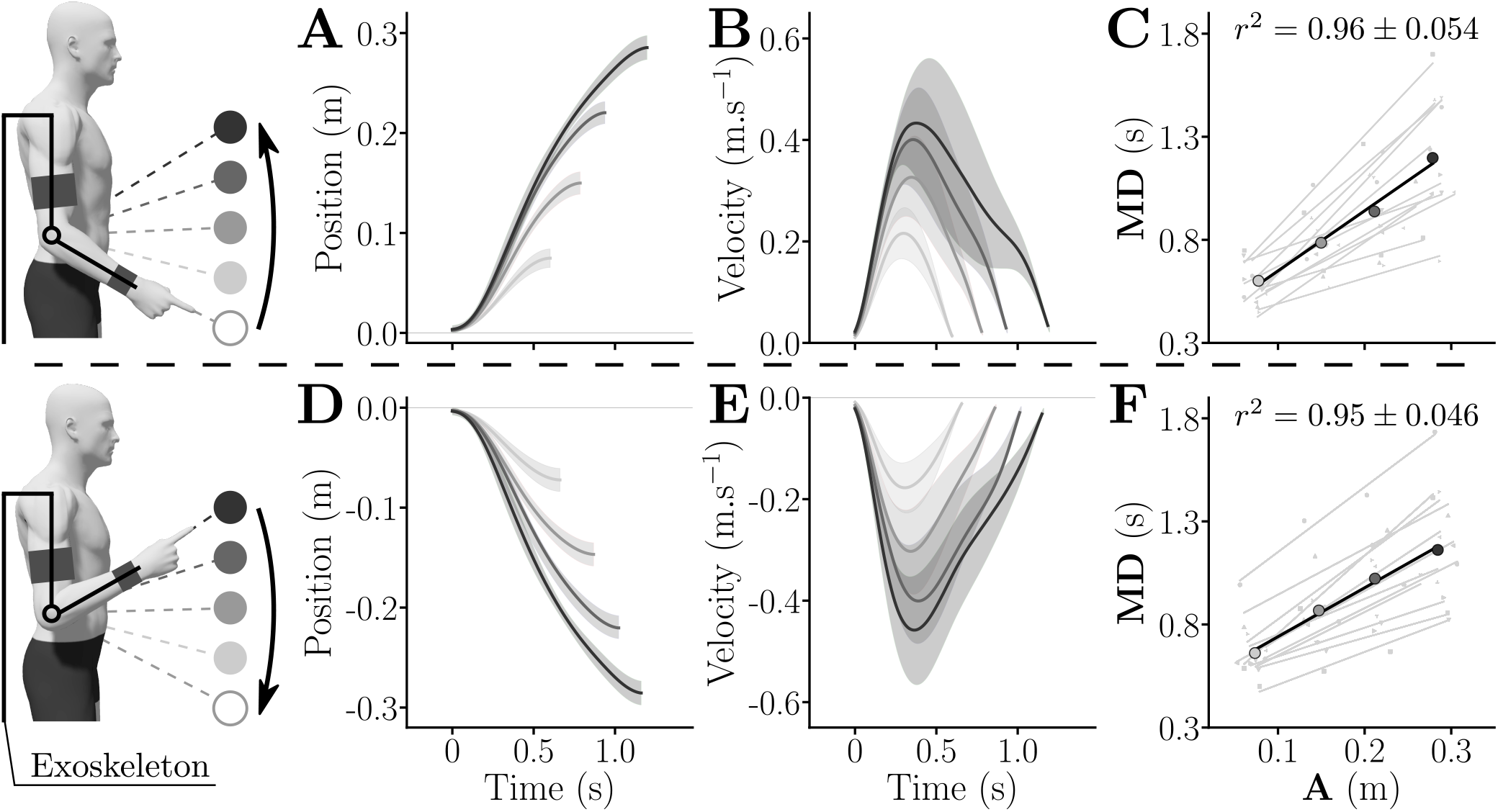
General kinematics averaged across all participants in the transparent exoskeleton for both upward and downward movements. **A,D.** Averaged positions for upward (**A**) and downward (**D**) movements across population. Standard deviations are depicted as shaded areas. **B,E.** Averaged velocities for upward (**B**) and downward (**E**) movements across population. Standard deviations are depicted as shaded areas. **C,F.** Amplitude-movement duration linear regressions for each participant (grey) for upward (**C**) and downward movements (**F**). The averaged behavior of the population is given in black. The average and standard deviation of the correlation coefficient across the population are given on their respective graphs.

The average affine fits across participants for upward and downward movements (black lines in Fig. 2C,F), which were used to compute the vigor scores with respect to the population average for each participant and each direction (see Equation 4), were as follows:

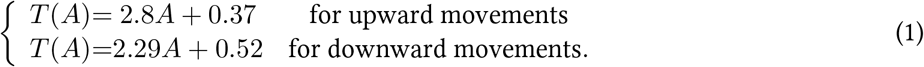

The spreading of individual vigor scores was shown to follow the same trend as in previous studies [32, 33], which was verified both for upward and downward movements (see Figures 3A,B). Moreover, the vigor scores of participants exhibited a strong consistency across directions (*r* = 0.89, *p* < 10^-3^, Fig. 3C). This analysis justifies *a posteriori* the use of the average amplitude-duration relationship of each participant to design the subject-specific assistive control law of the test session.

**Figure 3:**
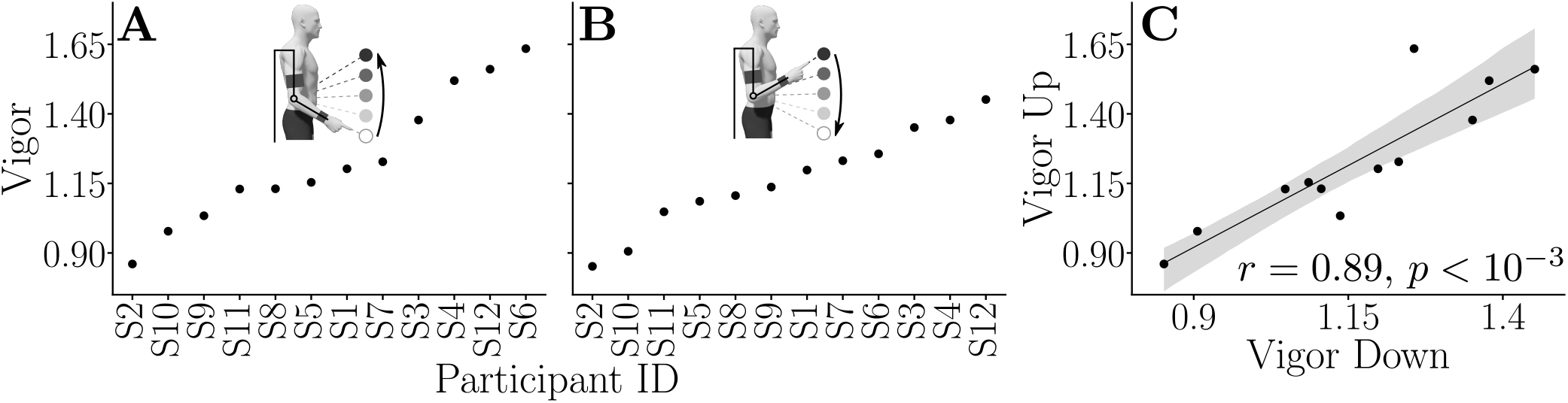
Individual vigor scores and consistency between upward and downward directions. **A.** Individual vigor scores for upward movements, sorted from lowest to highest. **B.** Individual vigor scores for downward movements, sorted from lowest to highest. **C.** Correlation analysis showing the consistency of vigor scores with regard to movement direction (Pearson correlation test).

### 2.2 Test session

In the test session, two amplitudes (17.5°, small amplitude (SA); 35°, large amplitude (LA)) and four assistance durations (*T_j_* = 100%*T*_*h*,0_, 200%*T*_*h*,0_, 400%*T*_*h*,0_ and 600%*T*_*h*,0_) were considered. The assistance was self-triggered by pressing a button with the left hand such that the participant could easily synchronize with the exoskeleton at the beginning of each movement. The assistance followed a minimum jerk velocity profile (see Equation 2 and [47, 48]). For upward movements, the planned trajectory was accurately followed if the participants remained inactive. For downward movements, the planned trajectory was followed only if the participants accompanied the robot’s movement by carrying their weight. Importantly, the participants could actively pull (*τ_h_* > 0) or actively push (*τ_h_* < 0) on the exoskeleton at any time during the movement. When they outpaced the planned trajectory, the exoskeleton applied a resistance proportional to the difference between the minimum jerk velocity and the actual velocity (see Equation 3). This resistance was calibrated on the basis of the MVF of the participant. It is worth noting that no resistance was applied to the participant if the actual velocity profile corresponded to the minimum jerk profile. Moreover, independently of the participant’s behavior, the exoskeleton was position-programmed near the target to remove any possible confound between minimizing time or preserving accuracy [37, 41, 49]. The experimental data were eventually compared to optimal control simulations according to the MTE theory, with the cost of time identified in the baseline session. We also compared these results to fixed-time simulations performed with the preferred duration of the average participant (*T*_*h*,0_) and with the assistance planned duration (*T_j_*), which can be seen as two extreme non-MTE strategies. The reader is deferred to the Materials and Methods for details.

Qualitatively, the experimental results indicated that the participants systematically saved time compared to the planned duration of the assistance (see velocity profiles in Figure 4 for LA and supplementary Figure S.1 for SA). Overall, these velocity profiles exhibited one main acceleration and one main deceleration even though they were less smooth than minimum jerk velocity profiles due to interaction with the robot. Peak velocities were larger than those of the assistance and movement durations were shorter. Noticeably, the MTE simulations were generally better at predicting the observed velocity profiles than simulations performed in fixed duration *T_j_* or *T*_*h*,0_.

**Figure 4:**
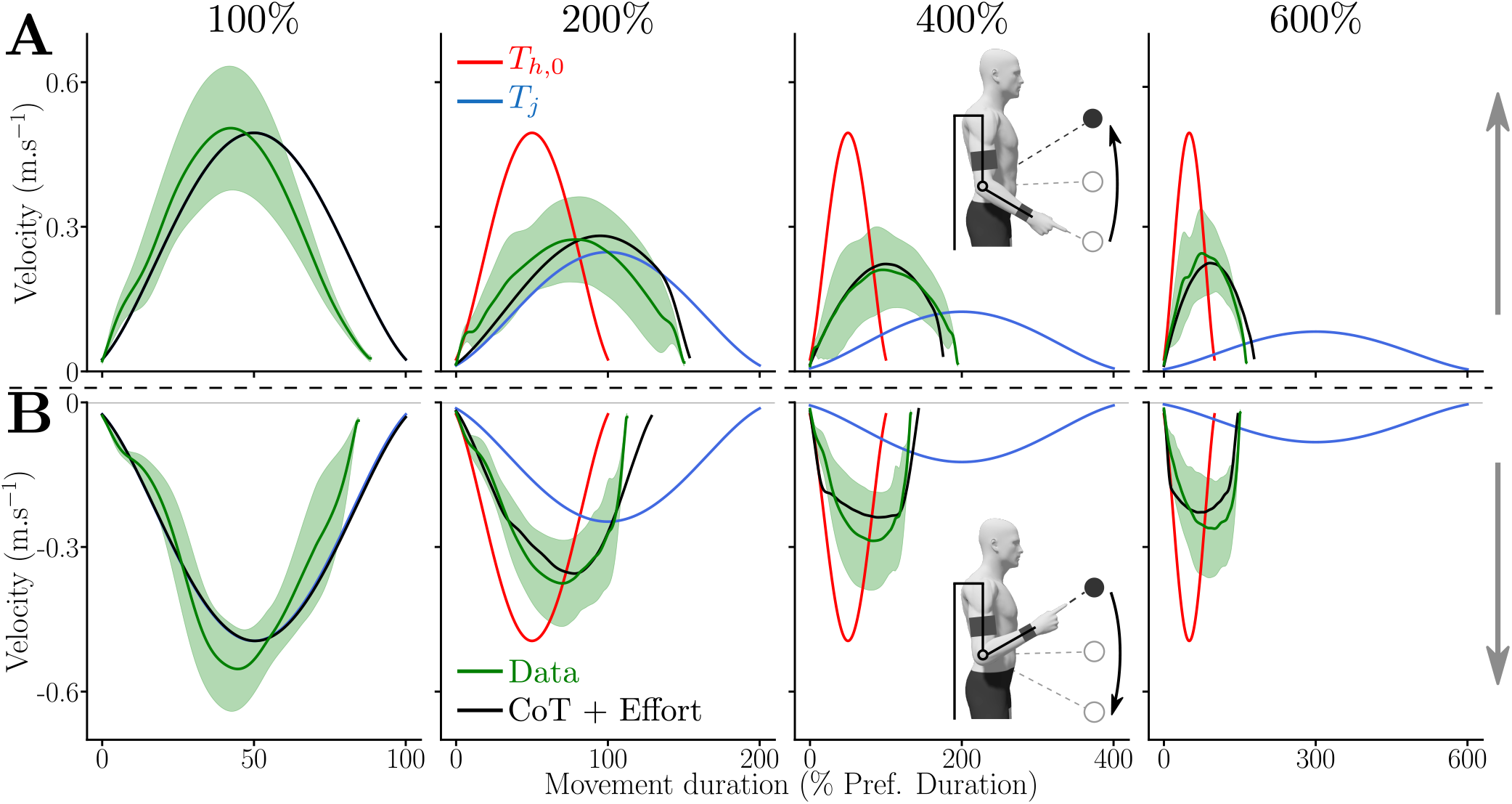
Average velocity profiles measured for the large amplitude (LA) and for each assistance duration. In green the average recorded velocity profiles and their standard deviation as a green shaded area, in black the optimal time-effort compromise, in blue the minimum jerk planned by the assistance and in red the constant time strategy. **A**. Upward movements. **B**. Downward movements.

Quantitatively, the participants’ behavior was described by three main parameters in this task: 1) the movement duration relative to the preferred movement duration (MD), 2) the maximum interaction force between the participant and the exoskeleton in percentage of the MVF from the agonist muscle group (i.e., flexors when moving upwards and extensors when moving downwards) and 3) the work of the interaction force. The first two of these parameters are normalized by individual data in agreement with the design of the experiment. The work is used as an absolute estimation of the additional energy expended by the participant to modulate the execution of the task (and possibly save time).

#### Movement duration

The MD measured during the experiment for the different assistance conditions, directions and amplitudes is depicted in Figures 5A,B,D,E.

**Figure 5:**
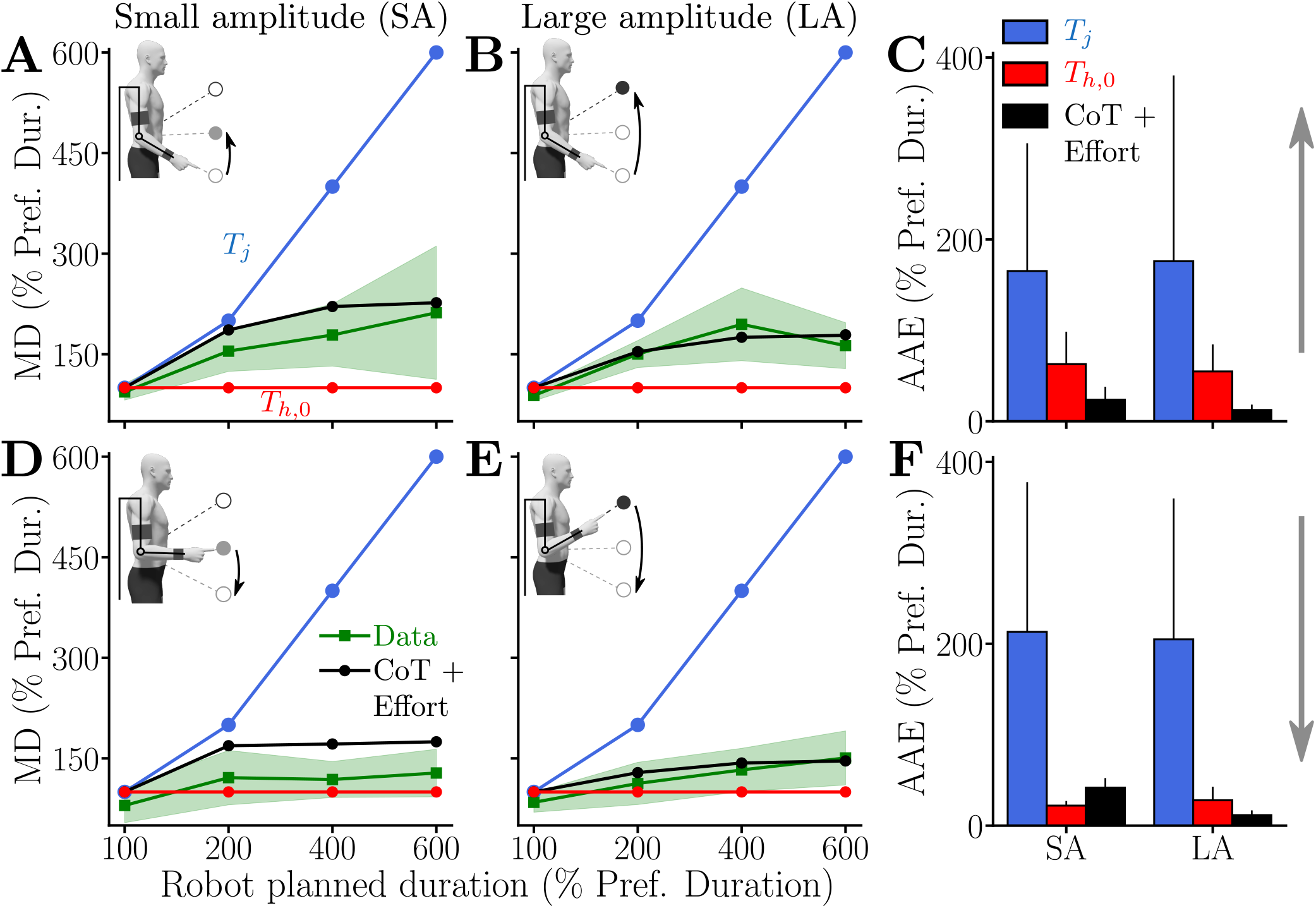
Chosen relative movement duration (MD) of participants when assisted by the exoskeleton with different *T_j_*. Average data are represented by green lines with standard deviation represented by green shaded areas. Outputs of different simulated motor strategies are depicted as follows: 1) in blue: simulation results with MD= *T_j_*, 2) in red: simulation results with MD= *T*_*h*,0_ and 3) in black: simulation results under a MTE hypothesis. **A,B.** Relative movement duration of upward movements for the small amplitude (SA, **A.**) and the large amplitude (LA, **B.**). **C.** Averaged absolute errors (AAE) of the different modeled strategies for both SA and LA for upward movements. **D,E.** Relative movement duration of downward movements for the small amplitude (SA, **D.**) and the large amplitude (LA, **E.**). **F.** Averaged absolute errors (AAE) of the different modeled strategies for both SA and LA for downward movements.

The results show that participants moved much faster than *T_j_* in the 200%, 400% and 600% conditions. This behavior was visible during movements of both SA and LA without any noticeable difference, and independently of movement direction (upward or downward). Nevertheless, participants did not return to their nominal MD in the task (*T*_*h*,0_, measured during the baseline session). Indeed, MD tended to increase as *T_j_* increased for both amplitudes and both directions. The increase in MD tended to be higher for upward than for downward movements. In the 100% condition, participants were on average slightly faster than during the baseline experiment, thereby suggesting that they were not completely passive and spent some effort to save even a little time.

These qualitative trends were confirmed by statistical Friedman tests. In particular, a main effect of the condition (*W* = 0.72, *Q*_3_ = 25.9, *p* < 10^-4^) and a main effect of the direction (*W* = 0.69, *Q*_1_ = 8.33, *p* = 0.0039)were observed. These tests also confirmed that movement amplitude has no effect on the normalized MD (*W* = 0.11, *Q*_1_ = 1.33 and *p* = 0.25).

Wilcoxon-Nemenyi pairwise comparisons were used as post-hoc tests to assess the most salient differences between conditions. First, upward movements of SA were significantly slower in the 200%, 400% and 600% conditions than in the 100% condition (in all cases: *p* ≤ 9.7 × 10^-5^, *D* ≥ 1.66 where *D* was Cohen’s *D*). The same trend was observed for downward movements of SA with movements performed in 200%, 400% and 600% conditions being significantly slower than in the 100% condition (in all cases: *p* ≤ 0.012, *D* ≥ 1.22). Second, upward movements of LA were significantly slower in the 200%, 400% and 600% conditions than in the 100% condition (in all cases: *p* ≤ 3.7 × 10^-5^, *D* ≥ 1.83). Upward movements of LA were also significantly slower in the 400% condition than in the 200% condition (*p* = 0.01, *D* = 1.08). Finally, downward movements of LA were shown to be significantly slower in the 200%, 400% and 600% conditions than in the 100% condition (in all cases: *p* ≤ 0.02, *D* ≥ 1.14) and those performed in the 600% condition were significantly slower than those performed in the 200% condition (*p* = 0.03, *D* = 1.05). In sum, these comparisons across conditions show that MD tended to increase as *T_j_* increased, independently of the direction and amplitude. Furthermore, comparisons were performed to analyze differences between upward and downward movements. Results were that MD was significantly lower for downward movements than for upward movements in LA in the 200% condition (*p* = 0.002, *D* = 1.43), in both SA and LA in the 400% condition (for both amplitudes: *p* ≤ 0.002, *D* ≥ 1.38) and only in SA for the 600% condition (*p* = 0.004, *D* = 1.12). In sum, upward movements were overall slower than downward movements in our task.

Overall, the MTE model replicated well the observed movement durations with the CoT identified during the baseline session. We evaluated the model predictions in terms of average absolute errors on MD (AAE, see Figure 5C,F). In agreement with the qualitative velocity profiles, the error of the MTE model was lower than those obtained when simulating movements with MD=*T_j_* (i.e., with the planned MD) or with MD=*T*_*h*,0_ (i.e. with the preferred MD of the average participant). The only notable exception was the AAE observed for downward movements in SA because the MTE prediction slightly overestimated movement duration in this condition.

#### Maximum interaction force

To understand the behavior of the participants in terms of effort, the maximum interaction force between the human and the exoskeleton relative to the MVF of the agonist group was analyzed (Figs. 6A,B,D,E). A positive value of this parameter means that the participant pulled on the exoskeleton (which is necessarily done actively) and a negative value means that the participant pushed on the exoskeleton (which can be done either passively –due to gravity– or actively).

**Figure 6:**
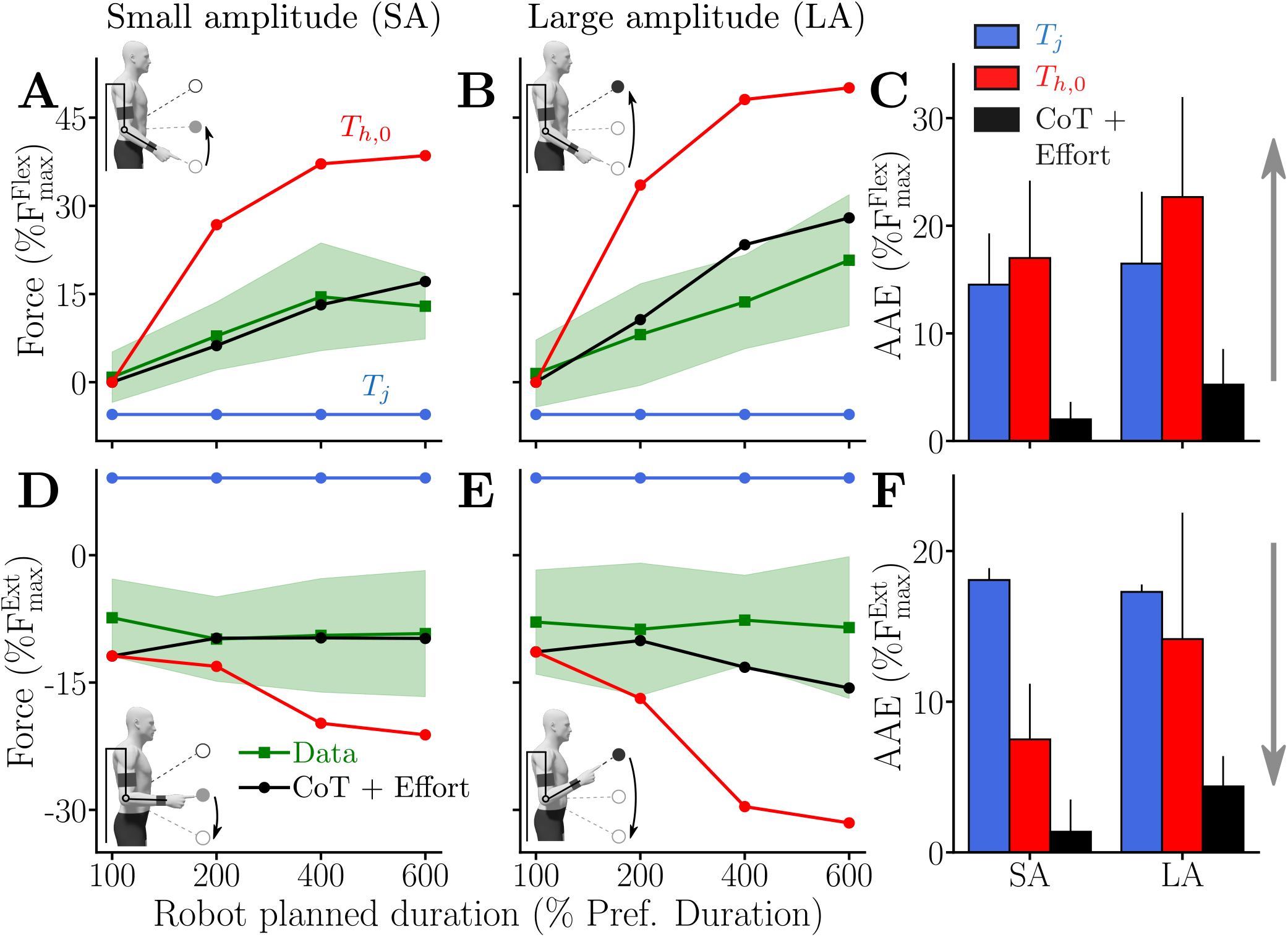
Maximum interaction force when the participant is assisted by the exoskeleton with different *T_j_*. Average data are represented by green lines and standard deviations as green shaded areas. Outputs of different simulated motor strategies (based on the dynamics from Equations 7 and 9, see Methods) are depicted as follows: 1) in blue: simulation results with MD= *T_j_*, 2) in red: simulation results with MD= *T*_*h*,0_ and 3) in black: simulation results under a MTE hypothesis. **A,B.** Maximum interaction force during upward movements for the small amplitude (SA, **A.**) and the large amplitude (LA, **B.**). **C.** Average absolute error (AAE) of the different modeled strategies for both SA and LA for upward movements. **D,E.** Maximum interaction force during downward movements for the small amplitude (SA, **D.**) and the large amplitude (LA, **E.**). **F.** Average absolute error (AAE) of the different modeled strategies for both SA and LA for downward movements.

The results show that, on average, participants tended to pull more and more on the exoskeleton as *T_j_* increased (Figs. 6A,B). Moreover, when moving upward in the 100% condition, the maximum interaction force between the participants and the exoskeleton was around zero on average. This means that participants tended to synchronize with the exoskeleton rather than being completely passive. Interestingly, their behavior was different during downward movements for which the maximum interaction force was globally constant and independent of *T_j_* (Figs. 6D,E). These trends were statistically confirmed by Friedman tests. In particular, a main effect of the assistance condition (*W* = 0.79, *Q*_3_ = 28.3, *p* < 10^-5^) and a main effect of the direction (*W* = 1, *Q*_1_ = 12, *p* < 10^-3^) were observed. Once again, movement amplitude did not seem to have a significant effect on the employed motor strategy, showing the robustness of the observations (*W* = 0.03, *Q*_1_ = 0.33 and *p* = 0.56).

Wilcoxon-Nemenyi pairwise comparisons on SA upward movements showed that participants applied significantly more force to pull the robot in the 200%, 400% and 600% conditions than in the 100% condition (in all cases: *p* ≤ 0.005, *D* ≥ 1.37). The participants also applied significantly more force to pull the robot in the 400% condition than in the 200% condition (*p* = 0.022, *D* = 0.87). On the contrary, no significant difference was found between the forces applied on the exoskeleton during downward movements. The same trends were observed during LA upward movements for which the participants applied significantly more force to pull the robot in the 400% and 600% conditions than in the 100% condition (in both cases: *p* ≤ 7.3 × 10^-4^, *D* ≥ 1.75). The participants also applied significantly more force to pull on the robot in the 600% condition than in the 200% condition (*p* = 0.0035, *D* = 1.27). As for SA, no significant difference was found between the forces applied on the exoskeleton during downward movements for LA. In summary, the participants applied an increasing maximal force on the exoskeleton as *T_j_* increased for upward movements. For downward movements, they applied a constant maximal force, independent of *T_j_*.

Furthermore, participants applied significantly different forces (in terms of absolute values) on the exoskeleton between upward and downward movements for all the conditions and for both SA (in all cases: *p* ≤ 2.46 × 10^-4^, *D* ≥ 1.85) and LA (in all cases: *p* ≤ 0.0011, *D* ≥ 1.58). Overall, the constant force applied when moving downwards (i.e., 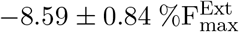) was remarkably close to the maximal effect of the weight of the human forearm and hand as estimated from anthropometric tables (i.e., 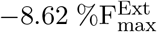). In summary, this suggests that participants were able to take advantage of gravity to save time when moving downwards.

Finally, we evaluated the model predictions in terms of maximum interaction force with the same error criterion as for MD (see Figure 6C,F). For this parameter, the MTE theory provided clearly the best results compared to alternative fixed-time strategies. On the one hand, simulations performed with MD=*T_j_* consistently resulted in a maximal interaction force whose sign was opposite to the measures. On the other hand, simulations performed with MD=*T*_*h*,0_ overestimated the interaction force that participants were apparently willing to use during the experiment. In contrast, the MTE theory correctly predicted the experimental trends across assistance durations, amplitudes and movement directions.

#### Work of interaction force

To get an absolute estimation of the total energy input (in Joules) from the participants onto the exoskeleton, we analyzed the work of the measured interaction force. A negative value for this parameter means that the interaction force mainly worked in the direction opposite to the motion. On the contrary, a positive value would reflect that the measured interaction force worked in the same direction as the motion. In particular, if a participant remains inactive during downward movements, this parameter should remain positive and approximately constant across assistance conditions for a given amplitude since the work of weight only depends on motion amplitude. The work of interaction force during the different experimental conditions is reported in Figures 7A,B,D,E.

**Figure 7:**
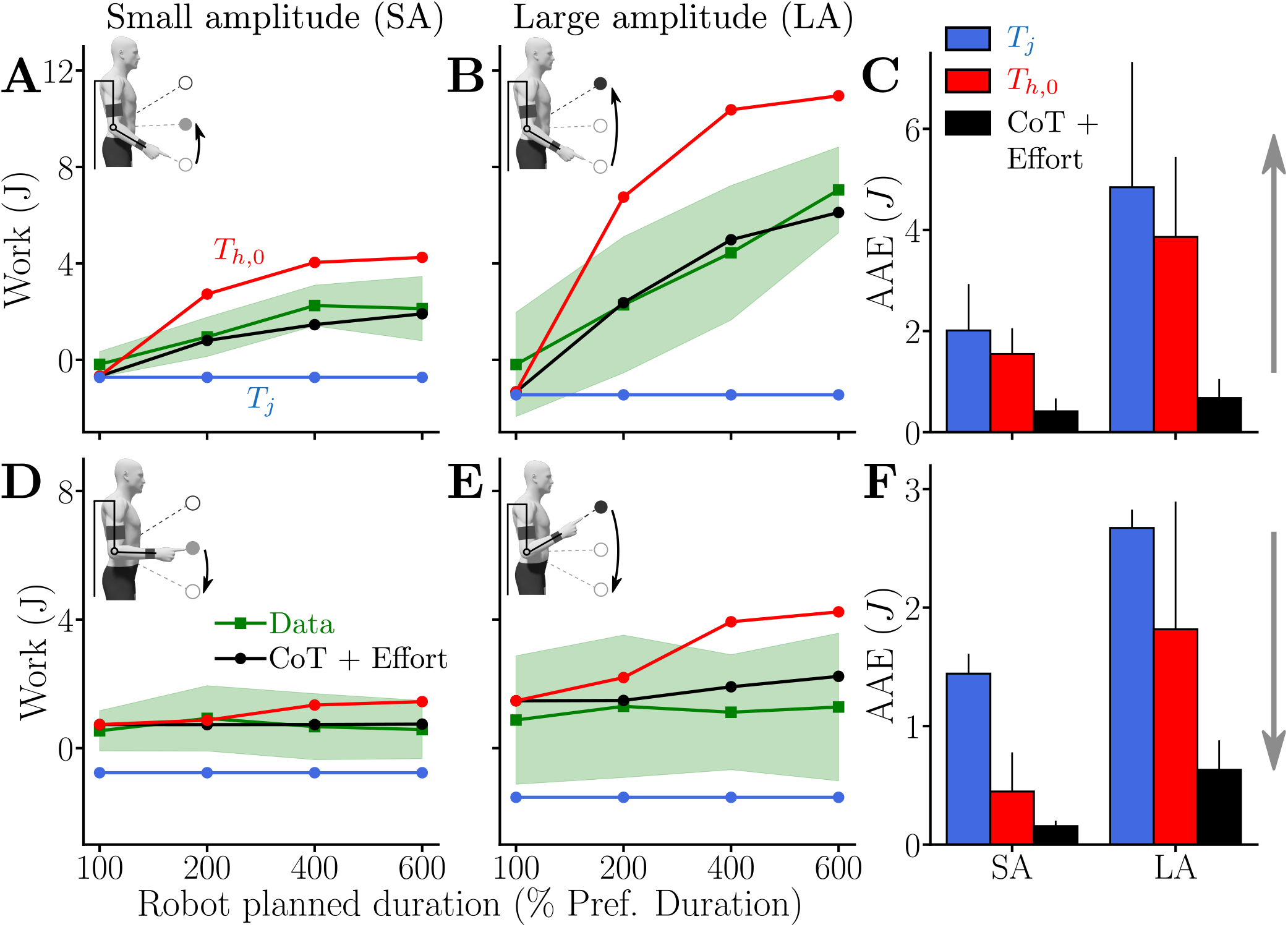
Work of the interaction force when the participant is assisted by the exoskeleton with different *T_j_*, average data are represented by green lines and standard deviations as green shaded areas. Outputs of different simulated motor strategies are depicted as follows: 1) in blue: simulation results with MD= *T_j_*, 2) in red: simulation results with MD= *T*_*h*,0_ and 3) in black: simulation results under a MTE hypothesis. **A,B.** Work for upward movements for the small amplitude (SA, **A.**) and the large amplitude (LA, **B.**). **C.** Average absolute error (AAE) of the different modeled strategies for both SA and LA for upward movements. **D,E.** Work for downward movements for the small amplitude (SA, **D.**) and the large amplitude (LA, **E.**). **F.** Average absolute error (AAE) of the different modeled strategies for both SA and LA for downward movements.

The average work in Joules turned out to be very similar to what was observed in terms of maximum interaction force. For upward movements, there was an increase in the human energy input to displace the robot when *T_j_* increased for both movement amplitudes. On the contrary, the work of interaction force was almost constant across conditions when moving downward. Overall, the energy input to the robot was higher for LA compared to SA movements, which was expected given the previous results on MD, maximum interaction force and the fact that work depends on the length of the trajectory. These trends were confirmed by Friedman tests that revealed significant main effects of assistance duration (*W* = 0.8, *Q*_3_ = 28.9, *p* ≤ 10^-5^), movement direction (*W* = 0.69, *Q*_1_ = 8.33, p = 0.004) and amplitude (*W* = 1, *Q*_1_ = 12, *p* ≤ 10^-3^). Since the main effect of movement amplitude could be expected for the work, the associated post-hoc tests will not be described hereafter.

Wilcoxon-Nemenyi pairwise comparisons revealed that, for upward movements in SA, the participants expended more energy in the 200%, 400% and 600% conditions than in the 100% condition (in all cases: *p* ≤ 7.31 × 10^-4^, *D* ≥ 1.66). Moreover, participants expended significantly more energy in the 400% and 600% conditions than in the 200% condition (in both cases: *p* ≤ 0.017, *D* ≥ 1.06).

The same trends were observed for upward movements for LA. Participants expended significantly more energy in the 200%, 400% and 600% conditions than in the 100% condition (in all cases: *p* ≤ 0.046, *D* ≥ 0.98). Furthermore, participants expended significantly more energy in the 600% condition than in the 200% and 400% conditions (in both cases: *p* ≤ 0.02, *D* ≥ 1.11). Furthermore, there was no significant effect of the assistance condition on the energy expended when moving downwards for both amplitudes. In sum, participants were willing to expend more and more energy as *T_j_* increased for upward movements. For downward movements, the work remained nearly constant, as did the maximum of the applied force.

The analyses conducted on the effect of direction revealed that the work of interaction force was higher for downward movements than for upward movements performed in SA in the 100% condition (*p* = 0.0086, D = 1.26). On the contrary, the work was always significantly higher for upward movements than for downward movements in the 400% condition (for both amplitudes: *p* ≤ 0.0051, *D* ≥ 1.42) and in the 600% condition (for both amplitudes: *p* ≤ 0.0051, *D* ≥ 1.36).

Interestingly, the nearly constant work of interaction force measured during downward movements (i.e., 0.68 ± 0.15 J for SA and 1.14 ± 0.17 J for LA) was remarkably close to the work of the human forearm’s weight for both amplitudes (*i.e.*, 0.71 J for SA and 1.42 J for LA using anthropometric tables [50]). This is in agreement with the previous observations made on the relative maximum force applied by the participants. Therefore, this result confirms that the participants took advantage of gravity-related efforts to accelerate the exoskeleton during downward movements, without actively producing work. Indeed, since the exoskeleton was controlled to never miss the target at the end of the motion, participants did not even have to expend energy to decelerate the system when approaching the target.

Finally, we evaluated the model predictions regarding the work of interaction force with the AAE, as for the other two parameters (Fig. 7C,F). Here again, the MTE theory provided the bests results in terms of AAE on work predictions. In particular, simulations performed with MD=*T_j_* consistently resulted in a negative work of interaction force, meaning that the simulated participant either actively pulled (i.e., *τ_h_* > 0 in these downward simulations) or passively pushed (i.e., the negative work is mainly due to weight in these upward simulations) against the exoskeleton. Furthermore, simulations performed with MD=*T*_*h*,0_ systematically overestimated the energy expenditure of the participants during the real experiment. In contrast, the MTE theory predicted well the work of interaction force across assistance durations *T_j_*, amplitudes and movement directions.

## 3 Discussion

In the present paper, we examined the extent to which participants rely on a common time-effort tradeoff under conditions that induce low or high energy costs to move with a certain vigor. To manipulate the usual relationship between vigor and effort, we used a robotic exoskeleton that could either assist or resist the participant’s motion. During upward movements, the results indicated that all participants saved time compared to the duration planned by the robotic assistance, thereby demonstrating a high propensity to expend energy to save time. During downward movements, a similar time saving was achieved by switching to a low effort strategy, thereby showing that participants did not mechanically associate saving time with expending more energy. Overall, the observed behavior was consistent with the minimization of a time-effort tradeoff.

Indeed, all participants consistently expended substantial amounts of energy to save time during upward movements but did not return to their nominal vigor in the task. The reason is likely that, when outpacing the reference trajectory of the robot, a viscous resistance was applied. Consequently, returning to the nominal vigor would have been admittedly possible but extremely expensive from an energetic point of view. For example, the work required to move with their nominal vigor would have been about 12 J per movement for the 600% and LA condition, Fig. 7B). Nevertheless, the energy expenditure consented by the participants remained high during upward movements, with an average work of 7.05 ± 1.78 J when pulling on the exoskeleton in this condition, which corresponds to an average work rate of 4.28 ± 2.08 J/s. For the sake of comparison, the work of the limb’s weight when performing an unconstrained elbow flexion of amplitude LA (accounting for most of the energy cost in these self-paced movements) was around 1.42 J, which amounts to an average work rate of 1.21 J/s with the mean vigor of our participants. Overall, these findings demonstrate that participants were willing to produce at least 3.6× their original work rate and spend about 5× their usual energy expenditure to get closer to their nominal vigor in the task.

This observation suggests that a cost growing quickly with time must be represented in the planning of such goal-directed actions. Otherwise, it seems difficult to explain why participants would expend so much energy to increase vigor of such point-to-point movements. Clearly, this additional human effort was not dedicated to control the final accuracy since it was always handled by the robot itself near the target. Moreover, participants started to energize the motion since its beginning. An alternative argument could be that participants just implemented a simple heuristic to solve the task at hand, without optimizing a genuine time-effort compromise. The rationale could be that it is a natural strategy because people are used to expend energy to produce movement. However, duration, interaction force and work systematically tended to increase with the robot’s planned duration during upward movements, which agrees with previous results obtained in an isometric task involving virtual movements [35]. The slower the assistance, the more participants pulled on the robot while consenting to reduce their vigor. This confirms that neither effort nor time were preserved or minimized alone across conditions. Interestingly, this energy expenditure pattern was very different for downward movements. Indeed, although MD followed a similar evolution, the energy expended by the participants was consistently very low across all assistance durations and significantly lower than for upward movements. Interestingly, the interaction measured in terms of force and work was indistinguishable from that of an inactive participant using only their weight to energize the motion planned by the exoskeleton. This capacity to exploit gravity is reminiscent of other results showing that the brain can optimally harness the effects of gravity to reduce effort during vertical arm movements [51–56].

Incidentally, this observation suggests that the strategy exhibited by participants during upward movements was not simply guided by a reluctance to inactivity. Nevertheless, in this task without explicit reward, it is unclear whether the hypothesized CoT only represents the temporal discounting of reward or not. Any type of cost growing with time could actually produce the same behavior. However, other authors have extensively studied how reward can affect movement vigor [14–19] and it is thus possible to assume that an implicit reward was associated with task achievement. By saving time on each trial, participants could leave the experiment earlier, which may be seen as a global reward as well. Since we did not explicitly manipulate reward in the task, we assumed that it was constant across conditions, which was reflected in our choice to use the same CoT in the model. Specifically, our paradigm modified the vigor-effort relationship by associating large or low effort costs to the nominal vigor of each participant in the task. This paradigm, together with the simulation results, provide evidence for the minimization of a common time-effort tradeoff across a wide range efforts, ranging from very active to mostly passive behaviors.

To derive our results, it is worth noting that we normalized the task to each nominal participant’s vigor and maximal voluntary force. Indeed, it is known that there is a large inter-individual variability on these parameters [16, 17, 30, 31, 33, 35]. Interestingly, we found no correlation between the maximum force and the nominal vigor in our participants (*R* = −0.12, *p* = 0.59). Without normalization, the results might have been more variable across participants in the test session. For instance, vigorous participants could have been more prone to expend significant amounts of energy to save time. However, what is considered a significant amount of energy may also depend on the strength of the participant. To avoid such complications, we opted for a normalization in terms of time and effort. Other analyses (not shown) revealed that the inter-individual differences were not consistent across conditions in the test session. Moreover, no correlation was found between the three main parameters under investigation and the nominal vigor of participants. Finally, one limitation of our study is that the conclusions were drawn from a relatively small number of participants. However, the statistical effect sizes were generally high (in most cases *D* > 1), meaning that our results reach a strong level of confidence. As expected, a post-hoc power analysis confirmed these conclusions by reaching a power of 0.93 for the smallest reported Cohen’s *D* (i.e., *D* = 0.98) and a power above 0.95 for all the other comparisons (i.e., with *D* ≥ 1.05).

Beyond that limitation, we believe that there are several interesting implications of the present results. In particular, with the emergence of new technologies for assisting human movement such as exoskeletons or co-bots, vigor may become a key factor to induce a more symbiotic interaction, whether it be for neurorehabilitation or for the prevention of musculoskeletal disorders at work [57–61]. Yet, current assistive robots can be relatively slow for safety concerns or computational reasons. This may cause unanticipated effects if, as predicted by the MTE theory, humans prefer to expend energy to save time when interacting with a too slow robot. The present study suggests that even a small reduction of vigor could lead the participants to attempt to strongly energize the motion if possible or reject the technology otherwise. Although the present paper does not allow to assess how the participants would actually behave during more complex tasks, for example involving more degrees of freedom or strong accuracy constraints, it still provides an interesting piece of information for the field of human-robot interaction.

Finally, understanding the invigoration of human movements is also essential for a better understanding of Parkinson’s disease, as underlined by several studies [62–65]. While bradykinesia is often associated with a misestimation of effort [62, 63], it could be equivalently explained by a misestimation of time [66]. One may speculate that the modulation of the basal ganglia’s input signals, which are known to determine movement vigor as a result of a dopamine/serotonin equilibrium [6, 8, 64, 65, 67–71], could regulate the interplay between time and effort via the direct and indirect pathways. Further analyses of the neural substrates involved in the time-effort tradeoff would help to clarify the mechanisms involved in action selection, in particular when it comes to set movement invigoration.

## 4 Materials and Methods

### 4.1 Participants and materials

#### Participants

A total of *N* = 12 participants (7 females) were involved in the experiment (mean age 28±6 years old, mean height 1.72 ± 0.07 m, mean weight 64 ± 12 kg, mean flexors MVF 236.5 ± 93.4 N and mean extensors MVF 173.4±67.4 N). All the participants were healthy, right-handed adults without known neurological disorder or injury that could have impacted the experiment. The participants gave their written informed consent as required by the Helsinki declaration to participate to the experiment, which was approved by the local ethical committee for research (CER-Paris-Saclay-2021-048).

#### MVF bench test

Individual MVF was measured on a custom H-shaped test bench made of aluminum profile and screwed into the ground to prevent any unwanted movement. A force transducer (SPEC) was mounted on the bench. This transducer was turned upwards for tests conducted on elbow extensors and downwards for elbow flexors.

#### Kinematics

Three-dimensional kinematics were measured by means of an optoelectronic motion capture device (10 Oqus 500+ infrared cameras, 100 Hz; Qualisys, Gothenburg, Sweden). The device tracked the position of twelve 10 mm reflective markers taped on the robot and seven 10 mm reflective markers taped on the participant. The markers taped on the participant were used to control the posture a posteriori. All the kinematic analyses were conducted on the recorded data of the marker taped at the end-effector of the robot. These analyses were equivalent to use the markers taped on the participant given that the position of each participant with respect to the exoskeleton was constant in the tested motion range [72].

#### Exoskeleton

The ABLE exoskeleton used in the experiment is an active upper-limb exoskeleton [73]. This exoskeleton was designed to be particularly compliant, which allowed to reach high levels of transparency [42, 74]. This exoskeleton replicates the three shoulder rotations (internal/external, adduction/abduction, flexion/extension) and the elbow flexion/extension of the human arm. The investigations here were restricted to the elbow joint of the exoskeleton for simplicity and the other joints were thus mechanically locked. Furthermore, the physical interfaces used to connect the human arm to the exoskeleton have been designed to maximize comfort and minimize unwanted interaction efforts [75, 76]. These developments were particularly important in the present context because the efforts transitioning at the level of the wrist interface could be intense, depending on the participant’s will to move fast.

#### Interaction efforts

A force-torque (FT) sensor (1010 Digital FT, ATI, maximum sample rate 7 kHz) was placed at the level of the wrist human-exoskeleton interface. This FT sensor could measure the six components (three forces and three torques) of the interaction efforts. During the present study, only the normal component of the interaction efforts was analyzed since it was the only one kinematically admissible by the human and exoskeleton elbow joints.

### 4.2 Experimental protocols

The baseline session was introduced to estimate the participants’ nominal vigor and their MVF. This was used to design the subject-specific assistive control law and identify the average cost of time of the participants in the task. The test session was introduced to assess the extent to which participants implemented a MTE when interacting with an assistive exoskeleton programmed to move at different speeds.

#### 4.2.1 Protocol of the baseline experiment

Before performing the pointing task with a transparent exoskeleton, the participants were asked to perform 6 trials of maximum isometric voluntary force (MVF) of 5 s each. Half of these trials were used to assess the MVF of the elbow flexors (mainly the biceps brachii and the brachioradialis) and the other half were used to assess the MVF of the elbow extensors (mainly the different heads of the triceps brachii). The participants pushed against a force transducer while their arm was vertical and their forearm horizontal. The contact between the participant and the force transducer was made of a foam-covered part to minimize discomfort and was located just behind the styloid process of the radius (flexors MVF tests) or the styloid process of the ulna (extensors MVF tests). The MVF was defined as the maximum force measured during the three tests.

Then, the participants were placed inside the exoskeleton and stood on a height-adjustable platform so that the position of the exoskeleton was always the same regardless of the height of the participant. They were asked to perform 32 flexions and 32 extensions of the elbow of an amplitude *A* ∈ {35°, 26.25°, 17.5°, 8.75°} (8 flexions and 8 extensions per amplitude) with the exoskeleton set in transparent mode (i.e. controller minimizing interaction efforts based on previous works [42, 43, 76]). Since only elbow flexions and extensions were required, the shoulder joints of the exoskeleton were mechanically locked. The target to reach to was defined as a green disk (4 cm diameter) displayed on a vertical screen and visual feedback of the current hand position was continuously displayed as a red disk cursor (1 cm diameter). The screen was placed at 1 m of the (fixed) elbow of the exoskeleton. The cursor position was updated in real-time to give a visual feedback of the current hand’s position, defined at the interaction between the line of the exoskeleton forearm segment and the plane defined by the screen. In all cases, the participants were instructed to execute those visually-guided movements at their preferred velocity. Throughout the movement, the target to reach was continuously displayed and it disappeared once the participant had stayed within it for 2 s with a velocity below 1 mm.s^-1^. The subsequent target was then displayed and so on, thereby alternating upward and downward movements.

#### 4.2.2 Protocol of the test session

In the test session, the participants performed a total of 4 blocks of 100 trials while the exoskeleton provided an assistance. Each block tested one of two amplitudes (i.e. *A* ∈ {35°, 17.5°}) with the same initial posture *q_i_*. Each block was divided in two sub-blocks of 25 trials testing different *T_j_*. The order of occurrence of the amplitudes and *T_j_* was pseudo-randomized across participants. Importantly, at the beginning of each sub-block, the participants were asked to relax using a message displayed on the screen for the first flexion and the first extension of each *T_j_*. This allowed to let the participant feel which movement was planned by the robot and the kind of assistance they could receive when remaining inactive.

The assistive control law was designed via a proportional-integral (PI) controller, the gains of which were set to allow the exoskeleton to track the reference trajectory in presence of the participant, when the switch to a viscous resistance was deactivated. The robot reference trajectory was derived from a minimum jerk model [47, 48]. This model is commonly accepted to generate smooth and bell-shaped velocity profiles. Despite known limits to capture velocity asymmetries observed due to gravity or accuracy [49, 77], this model was sufficient here to provide a human-like reference trajectory to be tracked by the PI controller. Precisely, the exoskeleton was controlled in position to minimize the tracking error *e* = *q_j_* – *q*, where *q* is the actual joint position of the robot and *q_j_* (i.e., the desired robot trajectory) is defined as follows:

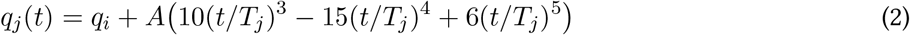

with *q_i_* the initial joint position of the robot, *T_j_* the robot’s movement duration determined after identification of the individual preferred duration *T_n_* for amplitude *A* (with *A* ∈ {35°, 17.5°})).

Once the assistance allowed the participant to reach to the target while remaining passive and without allowing the exoskeleton to switch its control mode, we considered the case where the participant could accelerate the motion, whether it be passively (with weight) or actively (meaning *τ_h_* ≠ 0). Since the gains of the PI controller were high enough to ensure a good tracking of the minimum jerk trajectory with the user inside the exoskeleton, the participant would not be able to significantly deviate from that trajectory without implementing an additional control mechanism. Therefore, to test our hypothesis, we introduced a criterion to detect when a participant overtook the robot and then switched to a viscous-like resistance while deactivating the PI controller.

The viscous-like torque resisting the human input was proportional to difference between the measured velocity 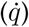 and the reference jerk velocity 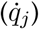. This viscous resistance was standardized according to the MVF of each participant, which resulted in the following expression:

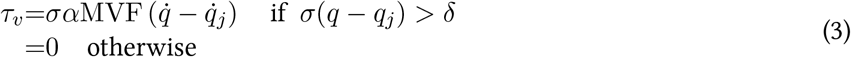

where *δ* = 0.02 rad is the deviation from the planned jerk trajectory in the direction of the movement (*σ* = 1 and *σ* = –1 for flexions and extensions respectively) and *α* = 0.1 is the resistance’s strength set to 10% of the MVF. The deviation *δ* was chosen so that weight was sufficient to outpace the exoskeleton for downward movements. Near the end of each movement, the robot was position controlled to ensure that the target was always accurately reached. This allowed to remove accuracy concerns for the participant and to minimize endpoint variance by design, thereby avoiding any unwanted speed-accuracy trade-off which could influence movement duration [37, 41, 49].

### 4.3 Data analysis

#### Kinematics

Three-dimensional position data of the marker placed on the exoskeleton’s end-effector were used to assess the movement kinematics. Position data from the other markers was used as control to monitor residual motions. Position data were filtered (low-pass Butterworth, 5 Hz cutoff, fifth-order, zero-phase distortion, *butter* function from the *scipy* package) as in previous studies [55, 72, 77]. Then, velocity and acceleration were obtained by numerical differentiation. Movements were segmented using a threshold set at 5% of the peak velocity of the considered movement.

For each participant, a vigor score (*vg_n_*) was computed following pre-existing methods based on movement durations [31, 35], as follows:

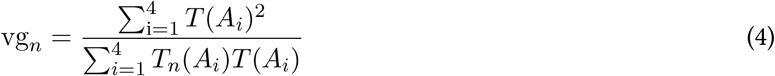

where *T*(*A_i_*) is the average duration computed from the population-based Equation 1 for amplitude *A_i_*, and *T_n_*(*A_i_*) is the averaged movement duration of the *n^th^* participant for amplitude *A_i_*. If the computed *vg_n_* is above 1, it means that the concerned participant moved overall faster than the population average. On the contrary, if the computed *vg_n_* is below 1, it means that the concerned participant moved overall slower than the population average.

#### Interaction efforts

As previously stated, the normal component of the interaction efforts was used to assess the force applied by the participants on the robot. These efforts were filtered (low-pass Butterworth, 5 Hz cutoff, fifth-order, zero-phase distortion, butter function from the scipy package) and segmented on the basis of the kinematic segmentation.

### 4.4 Statistical analysis

The statistical analyses were conducted using custom Python 3.8 scripts and the Pingouin package [78]. The normality (*Shapiro-Wilk* [79]) and sphericity (*Mauchly’s* [80]) of the data distribution were first verified. Since the results of these verification were not positive, Friedman tests were performed to check for possible main effects of the condition, the direction and the amplitude of movement. The significance level of the Friedman tests was set at *p* < 0.05.

Post-hoc comparisons were performed by means of non parametric pairwise Wilcoxon-Nemenyi comparisons. Their significance level was set at *p* < 0.05 and for each test the Cohen’s *D* was computed to analyze the effect size.

Finally, for information, a post-hoc power analysis was performed using the G*Power software (version 3.1.9.7, [81, 82]) in post-hoc mode with *α* = 0.05 and with the Cohen’s *D* reported in the paper.

### 4.5 Optimal control simulations

#### 4.5.1 CoT estimation

The CoT was identified on the basis of the averaged linear amplitude-duration relationship across all participants and directions (i.e. *T*(*A*) = 2.545A + 0.445, *r*^2^ = 0.99). The following model of the interaction dynamics was used when the robot was controlled in transparent mode:

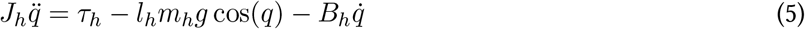

where *J_h_* = 0.043 kg.m^2^ was the human inertia, *m_h_* = 1.42 kg was the human forearm plus hand mass, *l_h_* = 0.17 m the distance between the elbow and the center of gravity of the forearm plus hand ensemble (these three population-average parameters were computed on the basis of anthropometric tables [50]) and *B_h_* = 0.05 Nm.s.rad^-1^ was the viscous coefficient of the elbow (this value was obtained in a previous study [83]). The joint position (respectively velocity and acceleration) was denoted by *q* (respectively 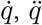). The assumption of perfect transparency was coherent with previous control developments [42, 43, 76], which allowed to cancel the significant effects of the exoskeleton on movement duration and peak velocity.

The minimum commanded torque change model was used in the present paper to predict human movement [84]. As a consequence, the state was defined as 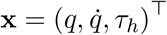 and the control variable was defined as 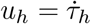. The cost function used to simulate movements from a starting state **x**_*i*_ = (*q_i_*, 0, *m_h_gl* cos(*q_i_*))^⊤^ to a final state *x_f_* = (*q_f_,* 0, *m_h_gl* cos(*q_f_*))^⊤^ in transparent mode and identify the CoT was as follows:

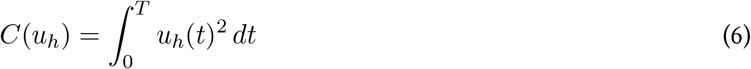

where *T* was estimated from the averaged amplitude-duration relationship for a given amplitude *A* = |*q_f_* – *q_i_*|. Then the procedure described by Equations S.1–S.3, based on the deterministic optimal control theory was applied to identify the CoT [29, 85]. After this procedure, our model was able to predict the nominal vigor of the average individual. Indeed, the addition of the CoT to the movement cost *C*(*u_h_*) yielded exactly the optimal duration corresponding to the experimental one for a movement joining **x**_*i*_ to **x**_*f*_.

#### 4.5.2 Simulations of possible behaviors with the assistance

Our experiment induced two main scenarios: one in which it was only possible to save time at the cost of an important energy expenditure (upward movements) and one in which being essentially inactive was sufficient to save time (downward movements). These two configurations were simulated separately given they suppose quite different interaction dynamics. Furthermore, each of these main configurations induced three possible scenarios: 1) actively pulling or pushing in the direction of the target (red shaded areas in Figures 1B,C), 2) remaining inactive (which is passively pushing, black dotted lines in Figures 1B,C) and 3) actively pushing or pulling in the opposite direction to the target (blue areas in Figures 1B,C). The latter scenario was unlikely from the MTE viewpoint and hardly doable in practice during upward movements because the assistance was performed by a relatively strong position control of the robot.

##### Prediction of human behavior when saving time is energetically expensive

First, the behavior of participants in a situation that did not allow saving time without expending energy was simulated (which corresponds to the red area in Figure 1B). This scenario was tested during upward movements with the jerk assistance in the present experiment. If the participant wanted to save time in this case, they needed to take control of both their own and the exoskeleton’s dynamics while counteracting the viscous resistance. The system dynamics was thus formulated as in Equation 7, and simulated from an initial state **x**_*i*_ = (*q_i_*, 0, (*l_h_m_h_* + *l_r_m_r_*)*g* cos(*q_i_*))^⊤^ to a final state **x**_*f*_ = (*q_f_*, 0, (*l_h_m_h_* + *l_r_m_r_*)*g* cos(*q_f_*))^⊤^.

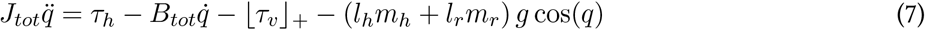

where *τ_h_* is the human torque, *J*_tot_ = *J_h_* + *J_r_* is the total inertia of the coupled system, *B_tot_* = *B_h_* + *B_r_* is the total viscous torque of the human and exoskeleton elbows respectively and (*l_h_m_h_* + *l_r_ m_r_*) is the total mass-length product inducing gravity related torques. The values of human parameters were the same as in Equation 5. The values of robot parameters were *J_r_* = 0.3 kg.m^2^, *B_r_* = 0.12 Nm.s.rad^-1^ and *l_r_m_r_* = 0.26 kg.m, which were identified following a preexisting procedure [42]. Finally, ⌊*τ_v_*⌋_+_ denotes that only the positive part of the viscous resistance is taken into account to prevent it from becoming an assistance at the end of the simulated movements (when 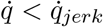, see the end of velocity profiles when *T_j_* ≠ 100%*T*_*h*,0_ in Figure 4A).

In the 100% condition simulations, participants tended to synchronize with the exoskeleton. Therefore, the torque applied by the assistance *τ_j_* was added to Equation 7. In the other conditions, this torque was not taken into account in the dynamics because participants systematically moved faster than the assistance, which deactivated it. Instead, the cost of following the assistance was computed separately (see blue vertical dotted line in Figure 1B for an illustration).

Finally, all these simulations were performed in free time (i.e. final time *T* ∈ (0, *T_j_*]) using an objective cost function that minimizes a compromise between time and effort as in Equation 6, using the previously identified CoT. This leads to an optimal movement time, illustrated by the black disk in Figure 1B). This cost function was as follows,

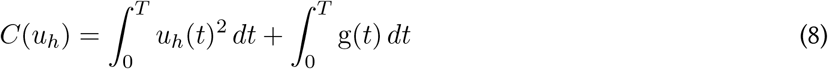

The MTE compromise computed with Equation 8 was then compared to the cost of following the assistance, which outputs are represented in blue in Figures 5–7, and to the cost of always moving at the preferred velocity, which outputs are represented in red in Figures 5–7.

##### Prediction of human behavior when saving time while being inactive is possible

Second, the behavior of participants when saving time was not necessarily energetically expensive was simulated (which corresponds to both the red area and black dotted line in Figure 1D). This case corresponded to downward movements with the jerk assistance in the present experiment. In this scenario, the weight of the participant and of the exoskeleton was helping to save time and naturally counterbalancing the viscous resistance. Moreover, the position control implemented at the beginning and end of movements allowed participants to be completely relieved of weight control if they wished to. In that case, only the inertia and natural viscosity of the human and robot segments and joints were handled by the participant. The system dynamics was thus simulated as in Equation 9, from an initial state **x**_*i*_ = (*q_i_*, 0, 0)^⊤^ to a final state **x**_*f*_ = (*q_f_*, 0, 0)^⊤^.

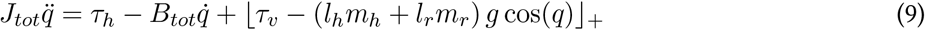

During simulations of downward movements, and contrary to those predicting upward movements, gravity related torques were directly compared to the viscous resistance and only positive values were taken into account in the dynamics. This simulated a natural compensation of all or a part of the viscous resistance by weight if participants pushed downwards or remained inactive (which corresponds to both the red area and black dotted line in Figure 1D). The simulations were then performed in free final time (i.e. *T* ∈ (0,*T_j_*]) using the same objective cost function as for upward movements (see Equation 8).

Finally, the case of participants pulling upwards in the opposite direction to the target was only simulated for a duration corresponding to *T_j_* as an illustration (represented in blue in Figures 5–7). Indeed, the cost of movement is trivially higher in that case given it induces an increase in both the cost of effort and the CoT (see dashed and dotted curves in the blue area in Figure 1D).

All the simulation parameters reported in the present paper were either direct results of the optimal control problem (relative movement duration) or computed using classical dynamics (interaction forces and work). All the simulations were performed using the Matlab (MathWorks) version of *gpops2* [86–88], which is a software based on an orthogonal collocation method relying on the *SNOPT* solver to solve the nonlinear programming problem [89].

## Acknowledgments

This work is supported by the French National Agency for Research (grant ANR-19-CE33-0009, EXOMAN project and grant ANR-22-CE37-0010, BasalCost project).

## Author contributions

Conceptualization: DV, OB, NV, BB.

Methodology: DV, BB.

Investigation: DV, GS, BB.

Data processing: DV.

Visualization: DV.

Supervision: OB, NV, BB.

Writing - original draft: DV.

Writing - review and editing: DV, OB, NV, BB.

## Data and materials availability

All data, code and materials used in the present study are available upon request to the corresponding author.

