## Supplementary for "The value of time in the invigoration of human movements when interacting with a robotic exoskeleton"

### 1 Average velocity profiles small amplitude

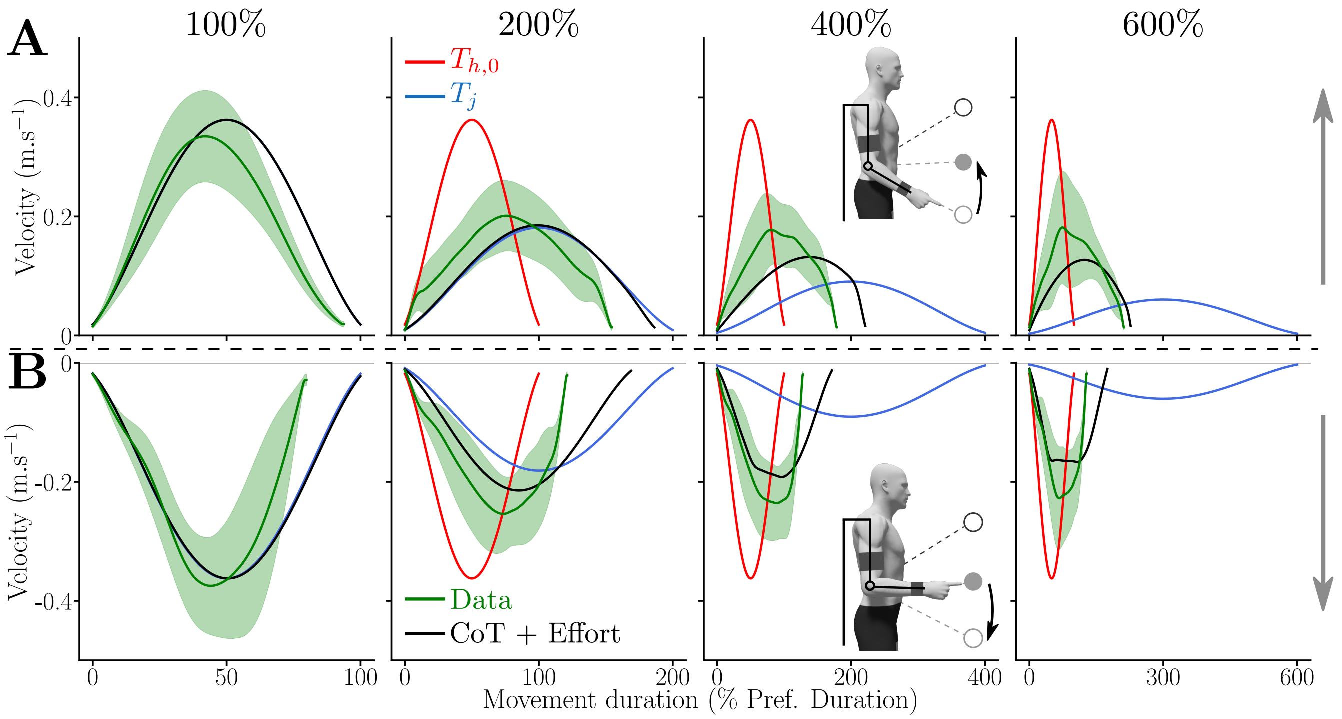


Figure S.1: Average velocity profiles measured for the SA and for each assistance duration. In green and standard deviation as a green shaded area, in black the optimal time-effort compromise, in blue the minimum jerk planned by the assistance and in red the constant time strategy. **A**. Upward movements. **B**. Downward movements.

### 2 Generalities regarding CoT estimation:

The objective was to assess the existence and quantify a possible effort/time optimal compromise. Therefore, simulations were designed in an optimal control framework with a composite cost function as in previous studies [1–4],

$$C(\mathbf{u}, t_f) = \int_0^{t_f} l(\mathbf{x}(t), \mathbf{u}(t))dt + \int_0^{t_f} \mathrm{g}(t)dt \tag{S.1}$$

where $\mathbf{u}$ was the control variable, $t_f$ was any final instant, $\mathbf{x}$ was the state vector, $l(\mathbf{x}, \mathbf{u})$ was the cost function representing effort and $\mathrm{g}(t)$ was the cost function associated with time (i.e. CoT).

In the present paper, CoT was estimated by simulating trajectories of various amplitudes in fixed time, based on average amplitudes and averaged amplitude/duration relationships measured in the the first part of the experiment pooling all upward and downward movements. In such simulations, CoT has been demonstrated to be the opposite of the Hamiltonian $\mathcal{H}_0$ associated with the optimal control problem [1],

$$\mathrm{g}(t_f) = -\mathcal{H}_0\left(\mathbf{x}(t_f), \mathbf{p}(t_f), \mathbf{u}(t_f), \lambda = 1\right) \tag{S.2}$$

where $\mathrm{g}(t_f)$ was the CoT for a final instant $t_f$, $\mathbf{x}$ was the optimal state vector, $\mathbf{p}$ was the only adjoint vector verifying the Pontryagin Maximum Principle with $\lambda = 1$ [5, 6]. Finally, $\mathbf{u}$ was the optimal control solving the problem. The Hamiltonian was then computed as follows,

$$\mathcal{H}_0\left(\mathbf{x}(t_f), \mathbf{p}(t_f), \mathbf{u}(t_f), \lambda = 1\right) = \mathbf{p}^T(t_f) f(\mathbf{x}(t_f), \mathbf{u}(t_f)) + \lambda l(\mathbf{x}(t_u), \mathbf{u}(t_f)) \tag{S.3}$$

where $f(\mathbf{x}, \mathbf{u}) = \dot{\mathbf{x}}$ represents the system dynamics, and $l(\mathbf{x}, \mathbf{u})$ is any non-negative cost function used in the motor control literature.
